# Robust Integration of Sparse Single-Cell Alternative Splicing and Gene Expression Data with SpliceVI

**DOI:** 10.1101/2025.11.26.690853

**Authors:** Smriti Vaidyanathan, Keren Isaev, Aaron Zweig, David A Knowles

## Abstract

Alternative splicing (AS) and gene expression (GE) are tightly related regulatory processes, critical for defining cell types and states, yet are rarely modeled together in single-cell analyses. This hinders a comprehensive understanding of cellular identity. We address this by introducing SpliceVI, adapted from MultiVI (Multi-modal Variational Inference) to specifically handle AS. Applied to a large multisample mouse Smart-seq2 dataset (*n* = 142, 315 cells/nuclei), SpliceVI jointly learns from both AS and GE using a partial variational autoencoder that effectively handles the sparsity and missingness of splicing data. We show that SpliceVI’s joint embeddings are more expressive and informative of biological correlates like age than a GE-only approach (scVI). SpliceVI also uncovers splicing-based differences between neuronal subclusters. This approach reveals the distinct yet synergistic relationship between AS and GE in shaping cellular diversity in mouse.

## 1 Main

Multi-modal profiling of individual cells offers detailed insights into cellular heterogeneity, function, and regulatory mechanisms. Technologies that simultaneously capture modalities such as gene expression (GE) and protein (e.g., CITE-seq [11]) or GE and chromatin accessibility (multiome assays [1]) are powerful for studying cellular programs. For example, analyzing paired RNA and accessibility in developing human brain organoids revealed a detailed “regulome” linking transcription factors to target genes in neurodevelopment and cell fate decisions [10].

To leverage the full spectrum of single-cell data, integration methods are essential for constructing unified representations from diverse contexts. Probabilistic frameworks like MultiVI [3] have demonstrated the efficacy of using variational autoencoders (VAE) to jointly model GE and chromatin accessibility. While various extensions exist, including models optimized for cross-modal translation (scButterfly [7]), imputation in partially matched datasets (JAMIE [8]), and scalable joint representation learning (multiDGD [20]), none have yet been adapted to the statistical challenges of alternative splicing (AS) data.

AS is a critical driver of cellular identity [13, 6, 14], yet it remains largely underexplored within modern multi-modal frameworks. This is partly due to data constraints: the ubiquitous 10x platform relies on 3’ or 5’ capture, failing to effectively sequence the internal regions required for splicing analysis. While Smart-seq2 protocols provide the necessary full-length coverage, AS quantification remains inherently sparse, as the total read count for a gene is partitioned across multiple competing splicing events, and has high missingness since AS cannot be quantified at all if the gene is not expressed or captured in a given cell. Existing methods like scQuint [4] have attempted to address this using VAEs, but treat AS as an isolated modality and rely on pre-imputation for missing data. Separate analysis of GE and AS variation masks the interdependence of transcriptional and post-transcriptional gene regulation, for example splicing factor abundance influencing target exon inclusion, or isoform-specific effects of TFs [15].

To address these limitations, we introduce SpliceVI, a novel deep generative model inspired by the MultiVI framework to incorporate both GE and AS information from single cells. We implemented SpliceVI as part of the scvi-tools codebase so that it seamlessly interoperates with the existing and widely utilized infrastructure [12]. SpliceVI uses a partial VAE architecture, enabling robust handling of sparse splicing events without requiring prior imputation for missing entries [17]. By explicitly modeling gene abundance and splicing variation within a unified latent space, and utilizing readily-interpretable linear decoders for both modalities, SpliceVI provides informative splicing imputation and captures meaningful substructure within well-defined cell types. We demonstrate that this integrated analysis yields a more granular view of cellular identity and begins to address the question of how much cellular diversity is driven by GE versus AS differences on an organismal level.

The SpliceVI architecture integrates these modalities using a partial VAE framework (Figure 1A, see Methods). This architecture is specifically engineered to handle the sparsity and missingness of AS data, employing a specialized encoder for observed PSI values and a Dirichlet-Multinomial (DM) likelihood [17] to accurately model splice junction counts (Methods). The entire model operates without requiring prior imputation. Having established this robust, missingness-aware architecture, we sought to test its performance and biological utility at scale. We applied SpliceVI to a large, paired mouse single-cell atlas (*N* = 142, 315 cells/nuclei) spanning diverse tissues and age groups (Isaev et al., manuscript in submission), benchmarking its performance against the gene-expression-only baseline, scVI [21] (Methods).

**Figure 1.**
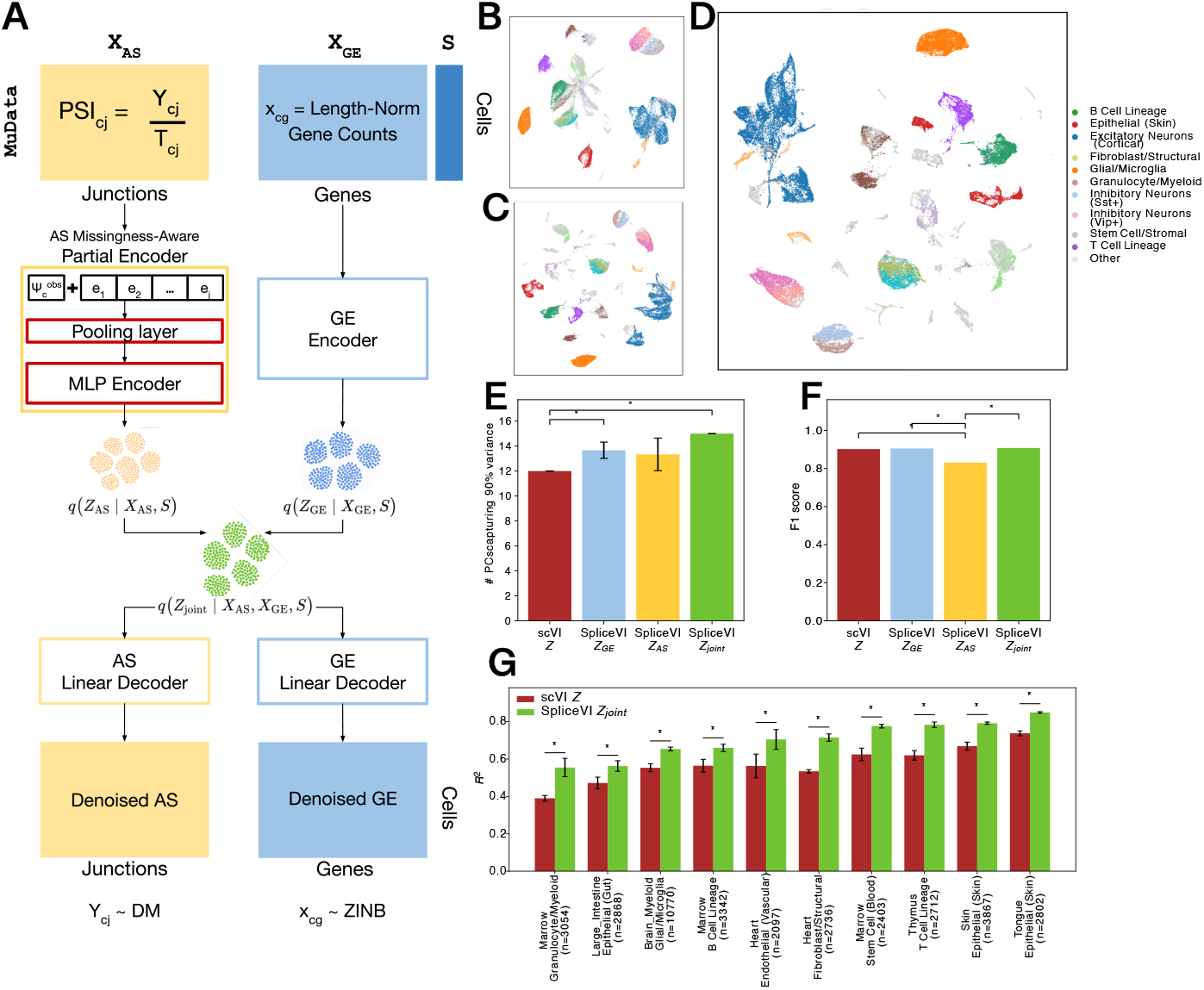
SpliceVI learns informative and expressive joint latent representations of AS and GE. **(A)**SpliceVI architecture: learnable junction embeddings *F*_*j*_ concatenated with PSI values, processed through a shared pooling layer, aggregated as 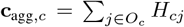, and fused with GE encoder outputs into joint latent space *Z*_*joint,c*_ for linear reconstruction. **(B)** UMAP of the AS encoder latent space *Z*_AS_. **(C)** UMAP of the GE encoder latent space *Z*_GE_. **(D)** UMAP of the joint latent space *Z*_Joint_. **(E)** Number of principal components required to explain 90% of the variance in latent spaces from scVI and SpliceVI (*Z, Z*_GE_, *Z*_AS_, and *Z*_Joint_) derived from the training data (n=99,620). **(F)** Average weighted F1 score of a multinomial logistic regression classifier trained on each latent space to predict GE-derived cell type labels in a held out test dataset (n=42,695). **(G)** Ridge regression performance for predicting mouse age in the ten tissue–cell type groups with the largest cell counts at 3, 18, and 24 months, using latent spaces from scVI and SpliceVI from the held out test dataset. Error bars for **(E-G)** show the 95% confidence interval of the corresponding score across n=3 replicate runs.

We found that the GE, AS, and joint latent spaces learned by SpliceVI exhibit clear structure, with each encoder capturing distinct cellular organization (Figure 1B-D). To quantify how much biological variation each space retains, we performed PCA on the latent representations from the AS encoder, GE encoder, joint space, and a single-modality (GE-only) baseline, linear scVI. We then measured how many principal components were required to explain 90% of the variance (Figure 1E). The joint latent space from SpliceVI was substantially more expressive than scVI’s latent space (∼14 PCs vs 12 PCs respectively), and the SpliceVI GE encoder also captured more variance across cells than scVI (15 PCs vs 12 PCs respectively). This increased expressiveness likely reflects the model’s training strategy, in which both decoders initially learn to reconstruct each modality from either latent representation (see Methods), encouraging each encoder to capture broad and complementary biological signals.

Further, to assess the biological relevance of the SpliceVI latent space, we evaluated how well it captures age-associated variation by training ridge regression models on each tissue–cell type group from single cells (n=26,596, 103 groups) in our held out test data. For each group, we compared the predictive performance using embeddings from SpliceVI (joint Z) versus scVI (GE only). Because AS varies across tissues and is known to exhibit age-related shifts [18], we expected the joint embedding to provide a stronger aging signal. Consistent with this, SpliceVI’s joint latent space achieved markedly higher *R*^2^ values across tissue-cell type pairs (Figure 1G). This suggests that AS, when jointly modeled with GE, captures non-redundant and biologically relevant variance that helps define age-associated cellular states.

To further assess biological interpretability, we compared the subcluster structure detected in the SpliceVI joint embedding with that obtained from the GE-only scVI baseline within cortical excitatory neurons (CENs). Using identical Leiden parameters, SpliceVI resolved six subclusters (Figure 2B), whereas scVI identified five (Figure 2A). To examine how these groupings differ, we performed pairwise differential expression and differential splicing (Methods) analyses for all subcluster pairs using normalized GE and DM-normalized PSI values from SpliceVI (Figure 2C-D). Several subcluster pairs defined by SpliceVI showed markedly higher numbers of significantly differentially spliced junctions than their scVI counterparts. The most pronounced contrast was between SpliceVI clusters CEN 3 and CEN 6, which displayed the highest number of DS junctions and DE genes than any pair identified using scVI labels.

**Figure 2.**
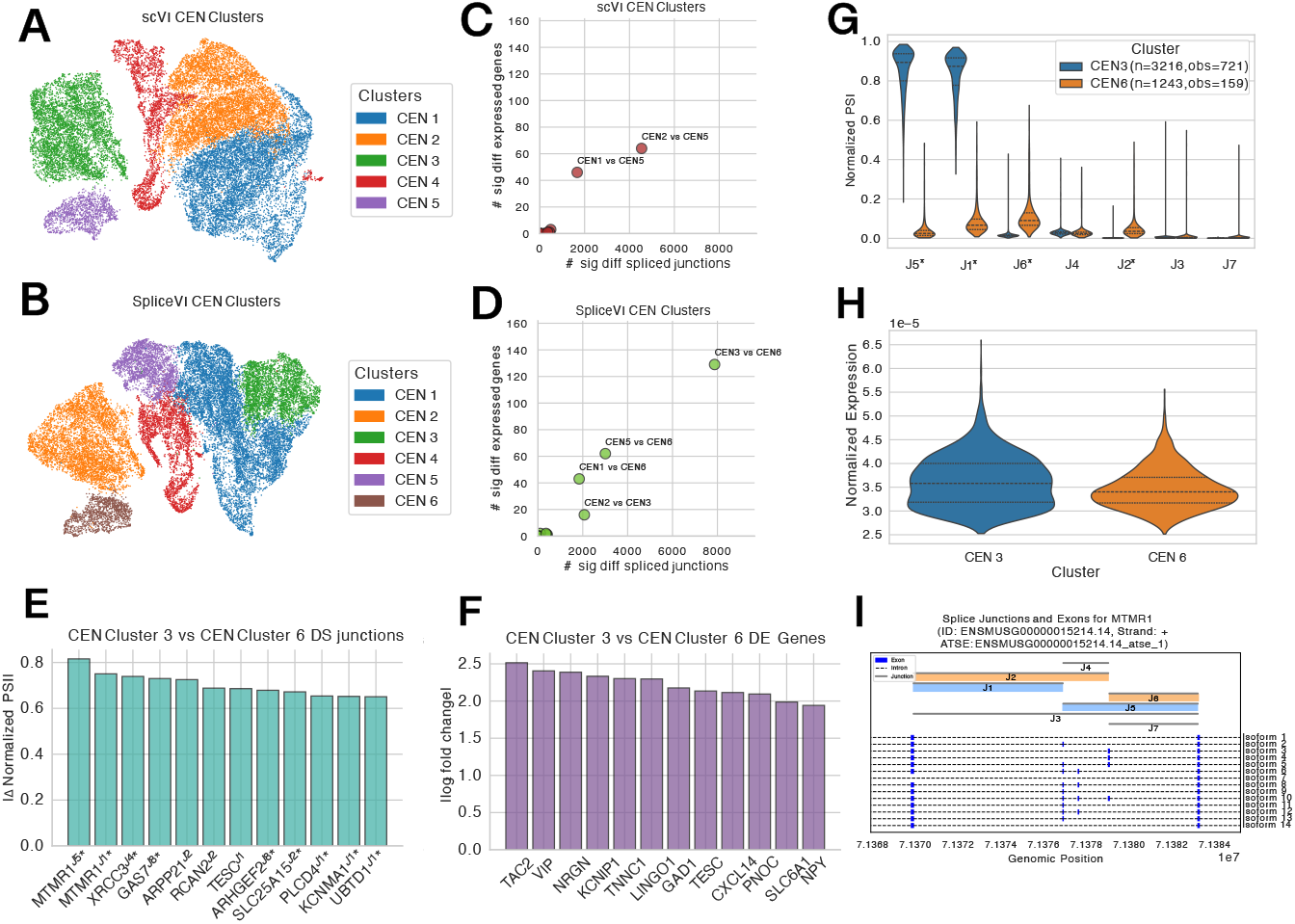
Cluster structure, differential analysis, and splicing characterization from SpliceVI. **(A)** UMAP of scVI latent space (*Z*) of Cortical Excitatory Neurons (CENs) with Leiden 0.1 clusters computed on the scVI latent representation. **(B)** UMAP of SpliceVI joint latent space (*Z*_Joint_) of CENs with Leiden 0.1 clusters computed on the latent representation. **(C, D)** Number of significantly differentially expressed genes and significantly differentially spliced junctions for all CEN Leiden cluster pairs, based on normalized GE and DM-normalized PSI; **(C)** uses clusters derived from scVI, and **(D)** uses clusters derived from SpliceVI. **(E)** Top ten differentially spliced junctions ranked by effect size (difference in DM-normalized PSI) between clusters CEN 3 and CEN 6, labeled by gene; junctions marked with ^*^ correspond to genes that are not significantly differentially expressed. **(F)** Top differentially expressed genes between CEN 3 and CEN 6, ranked by log fold change of normalized GE. **(G)** Violin plots of DM-normalized PSI for each junctions in ATSE *ENSMUSG00000015214*.*14_atse_1* within *MTMR1*, comparing clusters CEN 3 and CEN 6; junctions marked with ^*^ are significantly differentially spliced. In the legend, n is the number of cells in the cluster, and obs is the number of cells where that ATSE is detected. **(H)** Corresponding violin plot of normalized GE for *MTMR1* across the same CEN clusters. **(I)** Isoform structure of ATSE *ENSMUSG00000015214*.*14_atse_1*

To investigate this specific cluster pair, we visualized the top differentially spliced junctions and differentially expressed genes between CEN 3 and CEN 6 (Figure 2E-F). Most of the top DS junctions were not accompanied by DE of their parent genes, indicating that SpliceVI uncovers biological heterogeneity not detectable from expression alone. Notably, the two highest-effect splice junctions originated from the same Alternative Transcript Structure Event (ATSE, see Methods) in *MTMR1*, a gene that was not significantly differentially expressed. Within this ATSE, six junctions are present; four were identified as significantly differentially spliced. Examining the normalized PSI from each of these six junctions, cluster CEN 3 appears to prefer junctions J5 and J1, while CEN 6 favors J6 and J4 (Figure 2G). Normalized GE for *MTMR1* was comparable between the clusters, highlighting that this event is driven by AS rather than transcriptional differences (Figure 2H). An isoform diagram of the ATSE shows that these junctions correspond to mutually exclusive exon choices across most observed isoforms in a long read RNA-seq based reference, suggesting that the clusters differ primarily in isoform usage (Figure 2I). This pattern illustrates that SpliceVI identifies subtype-specific splicing programs that remain hidden in GE-only analyses.

Overall, SpliceVI demonstrates that joint modeling of GE and AS in single cells can reveal cellular heterogeneity not captured by either modality alone. By adapting a partial VAE architecture to handle sparse splicing data without prior imputation, SpliceVI effectively integrates these correlated transcriptional processes into a unified latent representation. Our results show that SpliceVI learns expressive joint embeddings that enable robust splicing imputation from GE, improve prediction of age-associated variation, and refine cellular subtype resolution by integrating both AS and GE, especially in neuronal populations where splicing diversity is functionally critical. The linear decoders facilitate interpretability, allowing future exploration of which latent features drive variation in each modality and how expression-splicing coordination shapes cellular identity.

Several extensions could further enhance SpliceVI’s utility and scope. Extending SpliceVI to operate directly on isoform-level quantification from long-read RNA sequencing would allow modeling full transcript abundance rather than splice junction-level signals alone [2]. The trained model could also be evaluated using transfer learning, mapping the learned embeddings onto human single-cell datasets to assess the conservation of AS and GE relationships across species.

## 2 Methods

### 2.1 Data

We processed publicly available datasets to generate inputs for the SpliceVI model training and evaluation (Figure 1A). The genome reference GENCODE vM19 (GRCm38) was used for all alignments and annotations. Two primary data sources were utilized to cover a wide range of tissues:

#### 1. Tabula Muris Senis (TMS)

Single-cell Smart-seq2 bam files from the Tabula Muris Senis consortium [22] were obtained from the Chan Zuckerberg Biohub AWS S3 bucket, accessible at s3://czb-tabula-muris-senis/.

#### 2. Allen Brain Atlas

Smart-seq2 based single-nuclei RNA-seq data (ssv4) from the Allen Institute for Brain Science [23] were obtained from the NCBI Gene Expression Omnibus (GEO accession: GSE185862) and the Allen Brain Map portal (https://portal.brain-map.org/atlases-and-data/rnaseq/mouse-whole-cortex-and-hippocampus-smart-seq). Processed count matrices, specifically exon-only counts, were used for GE analysis.

### 2.2 AS and GE Quantification

AS events were quantified from the Tabula Muris Senis (TMS) and Allen Brain (AB) Atlas Smart-seq2 data jointly using a detailed pipeline described previously (Isaev et al., manuscript in submission). This process aggregated 88,546 single cells from TMS and 53,769 single nuclei from AB (142,315 total single cells/nuclei) across diverse mouse tissues and ages, providing the largest paired single-cell AS/GE dataset of its kind. Briefly, splice junctions were extracted from single cell BAM files using Regtools [9]. To ensure complete and accurate splice site annotation, we leveraged a PacBio Long-Read Sequencing (LRS)-derived GTF annotation [19] to maximize the detection of known and novel junctions within the short-read data. Groups of overlapping junctions representing distinct local AS events were then clustered into Alternative Transcript Structure Events (ATSEs), analogous to LeafCutter clusters [16], using our `ATSEmapper` pipeline (https://github.com/daklab/LeafletFA-utils/tree/main/leafletfa_utils/atsemapper). An ATSE is defined as a set of two or more junctions sharing common splice sites.

The `ATSEmapper` pipeline yields the essential input matrices for SpliceVI: Junction counts (*Y*_*cj*_) for each splice junction *j* in cell *c* and the corresponding ATSE counts (*T*_*cj*_), which are calculated as the sum of all junction counts within that ATSE. From these raw counts, the Percent Spliced-In (PSI) value is derived using the ratio 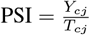, which serves as the core feature for the SpliceVI splicing encoder. This quantification was applied to our final mouse dataset, comprising a total of 89,831 splice junctions grouped into 28,015 ATSEs across 10,572 genes.

Raw gene count matrices were obtained from the TMS and AB datasets. For the AB Atlas data (single-nuclei RNA-seq), we utilized exonic reads only. We retained only common genes shared between the splicing input and GE that passed basic quality control filters. Gene read counts were normalized by average transcript length (due to the Smart-seq2 protocol) before inputting them into our model (*x*_*cg*_). Finally, the resulting GE and splicing modalities were combined into a single paired muData object.

#### 2.2.1 SpliceVI Input Data Structure

SpliceVI utilizes a MuData object that integrates GE and splicing modalities [5]. The GE modality contains transcript length–adjusted gene counts. The splicing modality contains three key data matrices corresponding to the quantification described above: splice junction counts, ATSE counts, and the PSI matrix. In addition, a binary mask layer within the splicing AnnData flags observed (1, non-zero ATSE count) versus missing (0, zero ATSE count) PSI values. This ensures that the partial encoder and reconstruction loss during training utilize only observed splicing events. We performed a 70–30 training–testing split on the full dataset (n=142,315) stratified on age and cell type. The training split was used as input for the model, which then performs an internal 90–10 split to obtain training and validation sets.

### 2.3 SpliceVI Model Definition

#### 2.3.1 Missingness-aware AS encoder

If the ATSE count for a given (cell, junction) pair is zero, the corresponding PSI value becomes 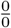, which is undefined (NaN) and should be treated as missing. In the context of a VAE, handling this in the decoder is straightforward, since these elements can simply be ignored and masked out when computing the likelihood. However, standard multi-layer perceptron (MLP) encoders require fully observed inputs. To overcome this limitation, SpliceVI adopts a *partial* VAE architecture for the splicing modality. This approach robustly models only the observed PSI values for each cell during training, circumventing the need for explicit pre-imputation of missing values, yet enabling imputation of all PSI values post-training. Our implementation draws inspiration from the EDDI framework [17], which provides a general strategy for VAE training from datasets with pervasive missing values using feature embeddings and exchangeable neural networks.

A core component of our splicing model is the junction-specific feature representation. We initialize learnable embeddings for each splice junction (*e*_*j*_ in Figure 1D) using Truncated Singular value decomposition (SVD). To obtain this initial embedding, PSI values across all cells are centered; this allows unobserved junctions in a given cell (missing PSI values) to be treated as zero for the purpose of the SVD dimensionality reduction only, minimizing the leakage of expression signal into the PCs. The first *D* (code_dim parameter in the model) principal components (loadings) derived from this centered PSI matrix serve as the initial values for these learnable junction embeddings.

The splicing partial encoder in SpliceVI processes only the observed splicing data for each cell *c*. Specifically, for each observed junction *j* in cell *c*:

1. The learnable junction embedding *e*_*j*_ ∈ ℝ^*D*^ is first *L*_2_-normalized and then scaled by the observed PSI value. This scaling step serves to weight the junction embedding by the magnitude of the observed splicing event.
2. The now normed and scaled learnable junction embedding *e*_*j*_ (of dimension *D*) is concatenated with the observed PSI value *ψ*_*cj*_.
3. This combined vector is passed through a neural network (an MLP), *h* which is shared across all junctions *j*, producing transformed features *H*_*cj*_.
4. These features *H*_*cj*_ are then aggregated (e.g., by summation or mean) across all junctions *j* observed in cell *c*, denoted *O*_*c*_, to yield an aggregated representation, 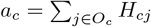.
5. This *a*_*c*_, optionally concatenated with relevant covariates (such as batch), is then processed by a cell-level MLP encoder to output the parameters (mean and variance) of the variational posterior for the splicing-specific latent variable *Z*_*AS*_.

#### 2.3.2 Decoder and Likelihood

For the GE modality, SpliceVI largely adopts the encoder architecture from the original MultiVI model [3]. This encoder processes transcript length adjusted gene counts (for Smart-seq2 data) for each cell to produce the parameters of a variational posterior for the GE-specific latent variable *Z*_*GE*_.

SpliceVI then integrates information from these two modalities. The modality-specific latent variables, *Z*_*AS*_ and *Z*_*GE*_, are combined to form a joint latent representation *Z*_*joint*_ for each cell. This integrated representation aims to capture the shared and distinct aspects of the cellular state reflected in both GE and AS.

Both modalities utilize log/logit-linear decoders to reconstruct their respective features from the joint latent representation *Z*_*joint*_, promoting interpretability. For splicing, the logit-linear decoder reconstructs the expected PSI *ρ*_*cj*_ for junction *j* in cell *c* from *Z*_*c*_ as,

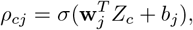

where **w**_*j*_ are the learnable weights and *b*_*j*_ are the learnable bias terms, and *σ*(·) is the sigmoid activation function. Similarly, for GE, a log-linear decoder maps *Z*_*c*_ to normalized GE *µ*_*c*_ and optionally the zero inflation logit(*π*_*cg*_).

This choice of log/logit-linear decoders for both modalities facilitates a more direct understanding of how latent features contribute to the observed molecular readouts.

##### Gene Expression

For each cell *c* and gene *g, x*_*cg*_ denotes the observed transcript length adjusted read count. By default we use a zero-inflated negative binomial (also the default in scVI),

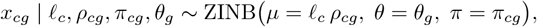

where *ℓ*_*c*_ is the cell’s size factor, *θ*_*g*_ the dispersion, and *ρ*_*cg*_ the expression prediction and *π*_*cg*_ the zero-inflation both from the decoder. Alternative likelihoods can be used and are identical to those in the original MultiVI implementation [3].

##### Alternative Splicing

For each cell *c* and junction *j, y*_*cj*_ is the observed read count. Let *a* be the set of junctions belonging to a specific ATSE, and define *T*_*ca*_ = ∑_*j*∈*a*_ *y*_*cj*_ as the count for the ATSE. We model,

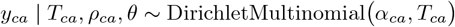

where *α*_*cj*_ = *ρ*_*cj*_*θ*_*a*_, with *ρ* the decoder output (PSI prediction) and *θ*_*a*_ a concentration parameter. In our implementation, *θ* for the Dirichlet-Multinomial is a learned per-ATSE concentration parameter. In both cases, we learn the logarithm of *θ* values as trainable parameters, initialized to log(100) with small Gaussian perturbations.

We also include options to use Beta-Binomial or Binomial likelihood, but generally recommend the DM.

##### Differential Splicing and Differential Expression Implementation

For GE, the normalized expression values employed in differential expression (DE) are simply the decoder’s predicted mean expression *µ*_*cg*_, rescaled by the size factor *ℓ*_*c*_ exactly as in scVI.

In contrast, for AS in differential splicing (DS), we do not use the raw decoder output *ρ*_*cj*_ directly. Instead, we compute a posterior-mean–stabilized PSI estimate that combines the decoder’s prediction with the observed junction counts for each ATSE. For a junction *j* belonging to ATSE *a*, the normalized PSI used in DS is

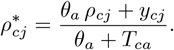

This is the posterior mean under the Dirichlet-Multinomial model, and yields a more stable estimate of junction inclusion, particularly for low-coverage junctions where *T*_*ca*_ is small.

We perform DE by applying the same methodology used by scVI and MultiVI for GE. For AS, the only difference is that the effect size is calculated using the difference in the predicted normalized PSI across both groups, instead of the difference in the logs (as is used for GE). For DE, we used a delta value of 0.25, and for splicing, 0.20; for both DE and DS, we used a FDR of 0.05.

#### 2.3.3 SpliceVI Model Implementation

SpliceVI was implemented by adapting the `MultiVI` and `MultiVAE` classes from a forked version of the scvi-tools codebase (v1.3.1; https://github.com/scverse/scvi-tools). We introduced new scvi-tools compatible classes, `SpliceVI` (model class) and `SpliceVAE` (module class), specifically adapted for RNA splicing inputs. These components replace the original data processing pipelines for chromatin accessibility with functionalities tailored to splicing analysis, while maintaining full compatibility with the existing scvi-tools API.

#### 2.3.4 SpliceVI Training Procedure

SpliceVI is trained end-to-end by maximizing an evidence lower bound (ELBO) that combines reconstruction losses for GE and AS, modality-specific KL terms to penalize deviation from a standard normal prior *N* (0, *I*), but no modality-alignment penalty (as is used in MultiVI).

To prevent posterior collapse and to encourage both encoders to learn meaningful structure, we employ a KL warmup phase over the first *n*_warmup_ epochs. During this period, the weight on each KL term is linearly increased from zero to one. In addition, we use a cross-modality routing strategy specifically during KL warmup to ensure that neither modality collapses. At each minibatch update, with equal probability, we randomly select one encoder (GE or splicing) and route its latent sample to *both* decoders. This forces the GE decoder to reconstruct GE from splicing-derived structure and the splicing decoder to reconstruct splicing from gene-expression-derived structure, stabilizing early training and improving identifiability of both encoders before the KL regularization reaches full strength.

We use a latent dimensionality of *K* = 25 for both encoders. The MLP in the AS Partial Encoder as well as the scVI GE Encoder both have a hidden dimension of 128 and have 2 layers. Both decoders are log/logit-linear. The junction-level embedding used in the Partial Encoder is size 32, with a h-layer hidden dimension of 64. We apply a dropout rate of 0.01 in the encoders, train with a minibatch size of 256, and optimize using AdamW with learning rate *η* = 10^−5^, weight decay 10^−3^, and gradient clipping at a maximum *ℓ*_2_-norm of 5. Models are trained for up to 500 epochs with early stopping based on validation ELBO, using a patience of 50 epochs, and the KL warmup length is set to 100 epochs. No batch correction key was used for the model results shown in this paper, although the model does have support for categorial and continuous covariates.

The model was trained on a single NVIDIA L40S GPU, taking approximately 8 and a half hours for 500 epochs on the full training dataset. The scVI benchmark was trained using the same dataset as SpliceVI with identical architecture wherever applicable and an identical training plan.

## 3 Acknowledgements

This work was supported by Columbia University and NYGC startup funds and NSF CAREER DBI2146398 to D.A.K. The content is solely the responsibility of the authors and does not necessarily represent the official views of the NSF.

## 4 Supplementary Methods

### 4.1 Inference Model

SpliceVI infers *ρ*_*cj*_ and *ρ*_*cg*_ from a latent variable *z* ∼ 𝒩 (*µ, σ*^2^) for each cell, integrated from modality-specific latent variables *Z*_*AS*_ and *Z*_*GE*_, which are learned with variational inference.

The posterior is approximated as:

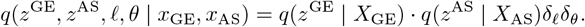

where the library size scale *ℓ* is estimated using a LibrarySizeEncoder, and per-gene dispersion *θ* is directly optimized, both inferred from the input data.

Modality-specific latent variables *z*_*GE*_ and *z*_*AS*_ are approximated as normal distributions with dimensionality *D*:

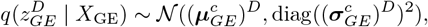

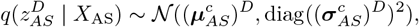

which are then combined as an average of the two, 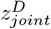.

### 4.2 Generative Model

The integrated *D*-dimensional latent representation *z*_*joint*_ is decoded by two modality-specific linear decoders: one for gene expression and one for splicing. These produce *ρ*_*cg*_ (normalized gene expression, dimensionality *n*, number of genes) and *ρ*_*cj*_ (normalized junction usage ratios, dimensionality *m*, number of junctions) for each cell *c*.

### 4.3 Objective Function

The model is trained by minimizing a loss function derived from the Evidence Lower Bound (ELBO). This loss combines a total reconstruction loss (ℒ_Total,*c*_), a KL divergence penalizing deviation of the joint latent variable *z*_*c*_ from its prior *p*(*z*_*c*_), and a KL divergence term to align the modality-specific latent representations.

The total reconstruction loss is the sum of losses for each modality:

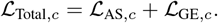

ℒ_AS,*c*_ is the negative log-likelihood for splicing (DM), and ℒ_GE,*c*_ is the negative log-likelihood for gene expression (ZINB).

The KL divergence KL(*q*(*z*_*c*_)∥*p*(*z*_*c*_)) regularizes the posterior distribution of the joint latent variable *q*(*z*_*c*_) towards a standard normal prior *p*(*z*_*c*_) = 𝒩 (0, 1):

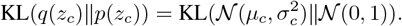

The total loss to be minimized is therefore:

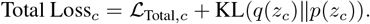

